# Calcium-triggered apoplastic ROS bursts balance gravity and mechanical signals to navigate soil

**DOI:** 10.1101/2025.01.07.631646

**Authors:** Ivan Kulich, Dmitrii Vladimirtsev, Marek Randuch, Shiqiang Gao, Matteo Citterico, Kai R. Konrad, Georg Nagel, Michael Wrzaczek, Jiri Friml

## Abstract

Reactive oxygen species (ROS) have been implicated repeatedly in multiple signaling processes in plants but the underlying mechanisms and roles remain enigmatic. Here, we developed live imaging of apoplastic ROS at the root surface. Different signals, including auxin, extracellular ATP and RALF1 peptide, all induce cytosolic calcium transients and apoplastic ROS bursts. Genetic and optogenetic manipulations identified calcium transients as necessary and sufficient for ROS bursts via activation of NADPH oxidases RBOHC and RBOHF. Apoplastic ROS bursts are not required but rather limit the gravity-induced root bending. Root bending is sensed by stretch-activated calcium channel MCA1 leading to NADPH oxidase activation at the stretched side. The resulting ROS production stiffens cell wall for better soil penetration. Apoplastic ROS thus provides a means to balance tissue flexibility and stiffness to efficiently navigate soil.

## Introduction

To colonize soil, plant roots grow downward but at the same time navigate through the complex environment of soil particles with varying densities. This involves dynamic changes in root growth integrating various cues, including the gravity, humidity, nutrients and pathogens (*1, 2*). Thus, roots need to combine flexibility for growth directions changes and stiffness for overcoming mechanical soil impedance. Research on petri dishes with homogenous media has led to detailed understanding of gravitropism, whereas the mechanisms underlying the tissue stiffening for the better soil penetrance remains enigmatic.

Root gravitropism involves accumulation of the plant hormone auxin at the lower side of the root elongation zone (*3*). Auxin there is recognized by the predominantly cytoplasmic receptor, AFB1, which mediates rapid, non-transcriptional responses such as cytosolic calcium (Ca^2+^) transients, decrease of membrane potential and cell wall alkalinization (*4*–*7*); all leading to root growth inhibition and downward bending.

When a growing root encounters an obstacle, it is first sensed in columella (*8*), resulting in ethylene production (*9, 10*). Afterwards, mechanical bending of the root is also perceived, causing Ca^2+^ signaling and alkalinization on the convex side of the root (*11*). The underlying mechanism and physiological role of these cellular processes are unknown.

Other cellular response linked to auxin is the apoplastic production of reactive oxygen species (ROS) (*12, 13*), which occurs also downstream of other signals including Rapid Alkalinization Factor (RALF1) peptide and its receptor FERONIA (*14, 15*), the extracellular ATP (*16*) (tissue damage indicator sensed by receptor kinase DORN1 (*17*)), or various other biotic and abiotic stresses (*11, 18, 19*). Apoplastic ROS are typically generated by NADPH oxidases but for different signals, different modes of upstream activation were proposed (*14, 20*–*26*); making the mechanistic dissection of its role challenging. Additionally, NADPH oxidases themselves are potent alkalinizing factors due to proton consumption during superoxide dismutation (*27, 28*). The ability to induce cell wall crosslinking and thus enhance cell wall rigidity (*29, 30*) makes apoplastic ROS an intriguing candidate for an additional, versatile growth regulation mechanism, nonetheless, this has not been established.

In this study, we employed a novel approach to live monitor ROS on the root surface. We identified cytosolic Ca^2+^ transients as a common mechanism for NADPH oxidase activation and the ROS bursts downstream of many signals. Genetic interference with ROS bursts revealed their essential role in cell wall fortification following mechanoperception, which is required for the soil impedance response and efficient soil penetration.

## Results

### Auxin triggers ROS burst by AFB1-CNGC14 signaling module

To monitor apoplastic ROS production, we measured the hydrogen peroxide (H_2_O_2_), which is a direct product of superoxide dismutation (*31*). The used Amplex RED stain predominantly localizes to the apoplast and exhibits red fluorescence in the presence of H_2_O_2_. We devised a novel method allowing a systematic and quantitative monitoring of apoplastic ROS. In our protocol, roots underwent two consecutive washes while being imaged: first with reduced 1 µM Amplex Red in mock media, followed by treatment with an elicitor added to fresh reduced Amplex Red. This setup enabled the measurement of fluorescent halos surrounding multiple root zones (Fig. 1A) and allowed the recording of even the initial seconds and dynamics of the ROS burst (fig. S1A,B); not accessible to previous ROS detection methods (*31*–*33*).

**Fig. 1.**
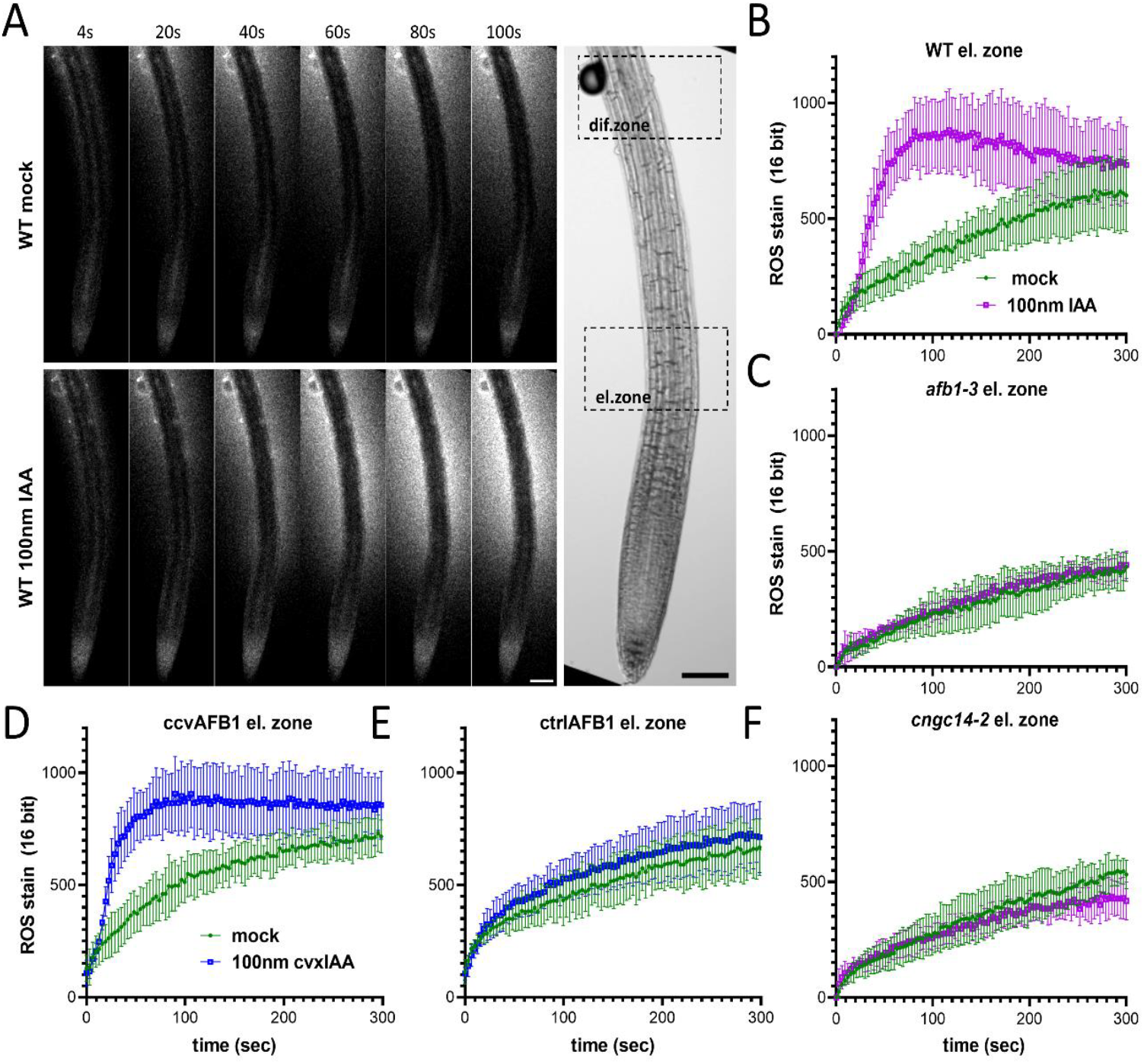
Apoplastic ROS production is activated by auxin by AFB1/CNGC14 module. **(A)** representative images depicting apoplastic ROS burst after IAA treatment as an Amplex red signal intensity. Differentiation and elongation zone (dif. zone and el. zone) depict regions where measurements were taken. Scalebar = 100µm. (**B** and **C**) IAA-induced ROS accumulation in the elongation zone of WT and *afb1-3* mutant. (**D** and **E**) 100nM cvxIAA-induced ROS accumulation in plants expressing ccvAFB1 and unmodified AFB1 (ctrlAFB1) under AFB1 promoter. (**F**) ROS accumulation of *cngc14-2* mutant elongation zone. All graphs represent average of >4 plants. Error bars are SD. Each experiment was conducted at least in two replicas.

Under mock conditions, ROS production was predominantly observed in the differentiation zone with a minimal contribution from the elongation zone (Fig. 1A,B; fig. S1C,D). Application of 100nM auxin (indole-3-acetic acid, IAA) resulted in only a slight signal increase in the differentiation zone but dramatically elevated ROS production in the elongation zone (Fig. 1A,B; fig. S1C,D), where the AFB1 cytoplasmic auxin receptor and the downstream component, the Cyclic Nucleotide Gated Channel 14 (CNGC14) are expressed (*34*). Similar results were achieved with 10nM IAA (fig. S1E). Notably, this ROS burst was absent in the *afb1-3*, which is impaired in the rapid auxin responses (*4*). To ascertain whether the AFB1-mediated IAA perception is sufficient to trigger the ROS burst, we investigated a synthetic receptor-ligand system, wherein concave AFB1 (ccvAFB1) exclusively binds to convex IAA (cvxIAA) but not unmodified IAA (*5, 35, 36*). In this system, cvxIAA induced ROS burst occurred exclusively in the presence of ccvAFB1 (Fig. 1D,E). AFB1 is thus necessary and sufficient auxin receptor mediating IAA-induced ROS burst. CNGC14 is another key component of rapid auxin signaling and gravitropism (*37*). Accordingly, in *cngc14-2* mutants, auxin-induced ROS production in the elongation zone was defective (Fig. 1F).

In conclusion, these observations identify apoplastic ROS burst as a new rapid, non-transcriptional auxin response. It depends on the dominantly cytosolic AFB1 receptor and the downstream CNGC14 Ca^2+^ channel.

### Multiple elicitors converge on calcium to trigger apoplastic ROS bursts

Given the essential role of the Ca^2+^ channel CNGC14 in the auxin-induced ROS burst and the well-documented involvement of Ca^2+^ in NADPH oxidase activation (*11, 38, 39*), we investigated the relationship between ROS production and Ca^2+^ homeostasis using Arabidopsis lines stably expressing intracellular Ca^2+^ sensor GCaMP3. Following 100nM IAA treatment, we observed a strong spatio-temporal coincidence between these responses with Ca^2+^ transients preceding the generation of apoplastic ROS (Fig. 2A,B; Movie S1) and the root elongation zone displaying the strongest intracellular transient and the most pronounced ROS burst.

**Fig. 2.**
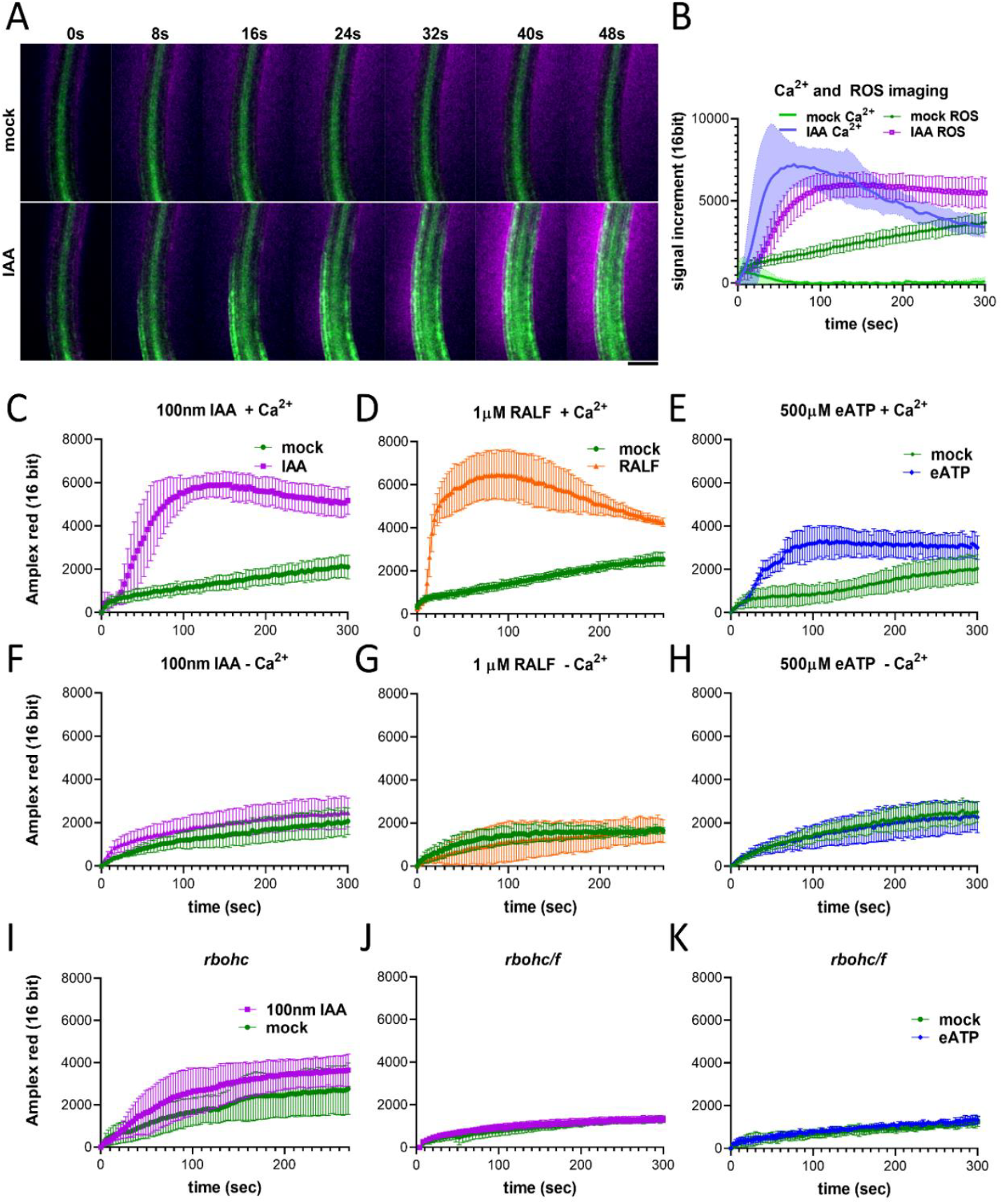
Apoplastic ROS bursts depend on the calcium signaling and NADPH oxidases RBOHC and RBOHF. (**A** and **B**) Apoplastic ROS burst (magenta) appears spatially and temporally after the intracellular transient (GCaMP sensor, green). Ca^2+^ transients elicited by IAA (**C**), RALF1 (**D**) and eATP (**E**) are not detectable without extracellular Ca^2+^ (**F** to **H**). (**I** to **K**) Genetic interference with NADPH oxidases; *rbohc* partially *and rbohc/f* fully abolishes ROS bursts induced by IAA or eATP. All graphs represent average of >4 plants. Errors represent SD. Each experiment was conducted at least in two replicas.

Subsequently, we examined the ROS burst of the root elongation zone in response to other known elicitors of intracellular Ca^2+^ transients, namely secreted peptide RALF1 (*40*) and extracellular ATP (eATP) (*41*). Akin to IAA, both RALF1 and eATP triggered apoplastic ROS bursts (Fig. 2C-E).

### Cytosolic calcium transients are necessary and sufficient to trigger apoplastic ROS bursts

To test the importance of Ca^2+^ transients for apoplastic ROS production, we prepared ½ MS media devoid of Ca^2+^ and supplemented it with either 3mM CaCl_2_ (+Ca) or MgCl_2_ (-Ca). Notably, in case of all tested ROS elicitors (IAA, RALF1, eATP), the absence of Ca^2+^ in the media abolished ROS burst (Fig. 2F-H).

To test whether Ca^2+^ is also a sufficient signal to induce apoplastic ROS, we used an optogenetic tool, the light-activated Ca^2+^ channel XXM2.0 (*42*). Blue light induced cytosolic Ca^2+^ transients robustly and repeatedly triggered apoplastic ROS bursts in roots (fig. S1F).

These observations revealed that Ca^2+^ transients downstream of various signals are necessary and sufficient for induction of the ROS bursts. This advocates for cytosolic Ca^2+^ as the universal intracellular mechanism mediating apoplastic ROS bursts.

### Root surface ROS bursts are fully dependent on RBOHC and RBOHF NADPH oxidases

To identify the source of apoplastic ROS burst, we focused on the NADPH oxidases. First, we used two known NADPH oxidase inhibitors VAS2870 and Diphenyleneiodonium chloride (DPI); which were, however, not sufficiently effective or specific, respectively (fig. S1F-K; Movie S2)

We then used the genetic approach, knocking out the NADPH oxidases with the highest expression in the epidermis of the root elongation zone, the RBOHC and RBOHF (*34*). Knockout mutant of *rbohc* (salk_016593C; (*43*)) showed visibly reduced, but still present ROS burst (Fig. 2I). Using CRISPR/CAS9, we obtained two lines with 300bp deletion at the 5’end of the *RBOHF* in the *rbohc* mutant background. The resulting *rbohc/f* mutants completely abolished ROS burst after IAA and eATP (Fig. 2J,K).

These observations identified RBOHC and RBOHF as two NADPH oxidase isoforms responsible for the production of majority of root surface ROS bursts downstream of multiple signals.

### Apoplastic ROS and pH regulation by NADPH oxidases in root gravitropism

*rbohc/f* mutants with undetectable apoplastic ROS bursts provided a genetic means to investigate their role in plant growth and development. Except for the root hair rupture phenotype (fig. S2A), *rbohc/f* roots exhibited normal growth in both, control conditions and when supplied with 10nM IAA (fig. S2A-B). This result was consistent also at pH 5.0, which enhances some *rbohc* phenotypes (*44*). Accordingly, the normal pH of the apoplastic space in the root elongation zone of *rbohc/f* mutant (fig. S2C) was detected using Fluorescein-5-(and-6)-Sulfonic Acid, Trisodium Salt (FS) (*45*). Furthermore, using microfluidic root chip, we observed normal rapid auxin-induced root growth inhibition in *rbohc/f* mutants (fig. S3D,E; Movie S3).

Normal root growth responses to auxin imply normal gravitropic bending of the *rbohc/f* mutant. Nonetheless, in contrast to the *cngc14-2* mutant, which bends visibly slower, its bending was slightly faster than in the wild type (WT) (fig. S3A). Faster bending of the *rbohc/f* mutant implies greater pH difference between the apoplasts of upper and lower elongation zone during gravitropic bending, which we confirmed by monitoring apoplastic pH using FS (fig. S3B,C). This was caused by more acidic upper side in *rbohc/f* mutants, while bottom alkaline halo remained as in WT (fig. S3B,C), again in contrast to the well-described *cngc14-2* mutant (*45*). These observations point towards importance of NADPH oxidase activity at the upper (convex) side of the root during gravitropic bending. Consistent with this, also Ca^2+^ transients and alkaline halo appear on the upper side of the root at later time points, when actual bending is in progress (fig. S3B,D).

In summary, NADPH oxidases and ROS production are not required for the auxin-induced rapid root growth inhibition and thus for root gravitropic bending. Instead, they limit the bending by contributing to the alkalinization on the upper (stretched) side of the root at later stages; a previously unobserved phenomenon, which might be related to mechanical stretching of root tissues during bending.

### MCA1-based mechanoperception induces apoplastic ROS for cell wall stiffening and soil penetration

To test, whether NADPH oxidases-generated apoplastic ROS bursts are related to mechanical stretching of tissues, we used a simple assay where we gently bend roots while imaging. Forced mechanical bending of the root resulted in a massive Ca^2+^ transient at the convex side of the root as previously reported (*11*). The Ca^2+^ transient was immediately followed by a ROS burst (Fig. 3A,B; Movie S4) and apoplast alkalinization (fig. S4). This shows that mechanical stretching induces Ca^2+^ transients, ROS burst and apoplast alkalinization.

**Fig. 3.**
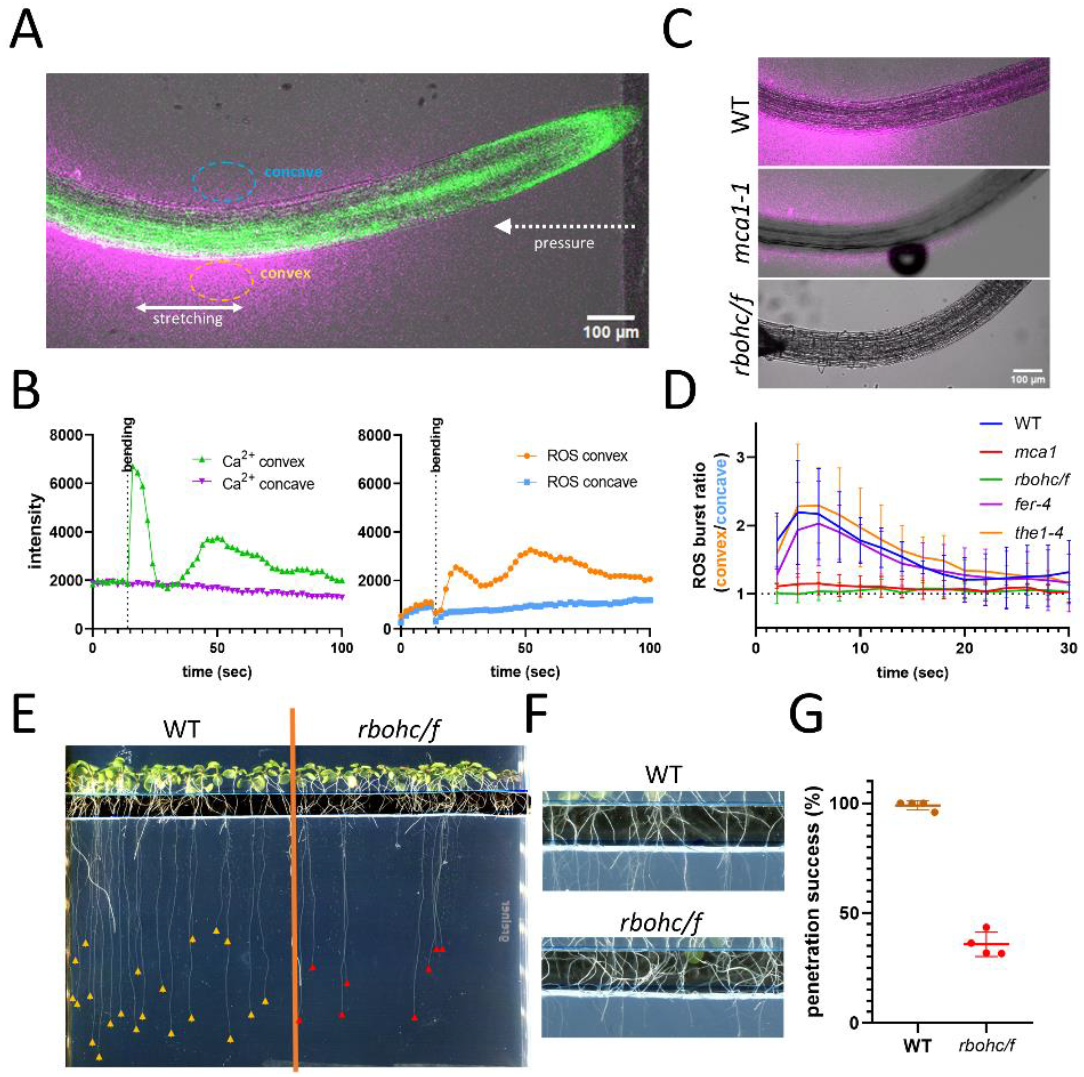
Mechanical tension induces NADPH oxidase dependent ROS burst via MCA1 and is required for the root-soil penetration. **(A)** Apoplastic ROS burst (magenta) and Ca^2+^ transient (GCaMP sensor, green) appear at the convex side after mechanical bending of the root. (**B**) quantification of representative bending event. Five similar replicates were obtained. (**C**) representative images of the ROS burst after mechanical bending of WT, *mca1-1* and *rbohc/f* mutants. (**D**) Quantification of the ROS bursts evoked by bending. Data represent ratio of the Amplex red intensity at the convex/concave sides. Average of 4-6 replicas with a single root. (**E**) representative image of the root-soil penetration experiment, where roots penetrate media solidified with 0.8% agar. Yellow and red arrows highlight the root tips. (**F**) detail of the air gap, which is filled by *rbohc/f* roots growing backwards. (**G**) *rbohc/f* penetration success calculated as % of shoots which penetrated the agar.

**Fig. 4.**
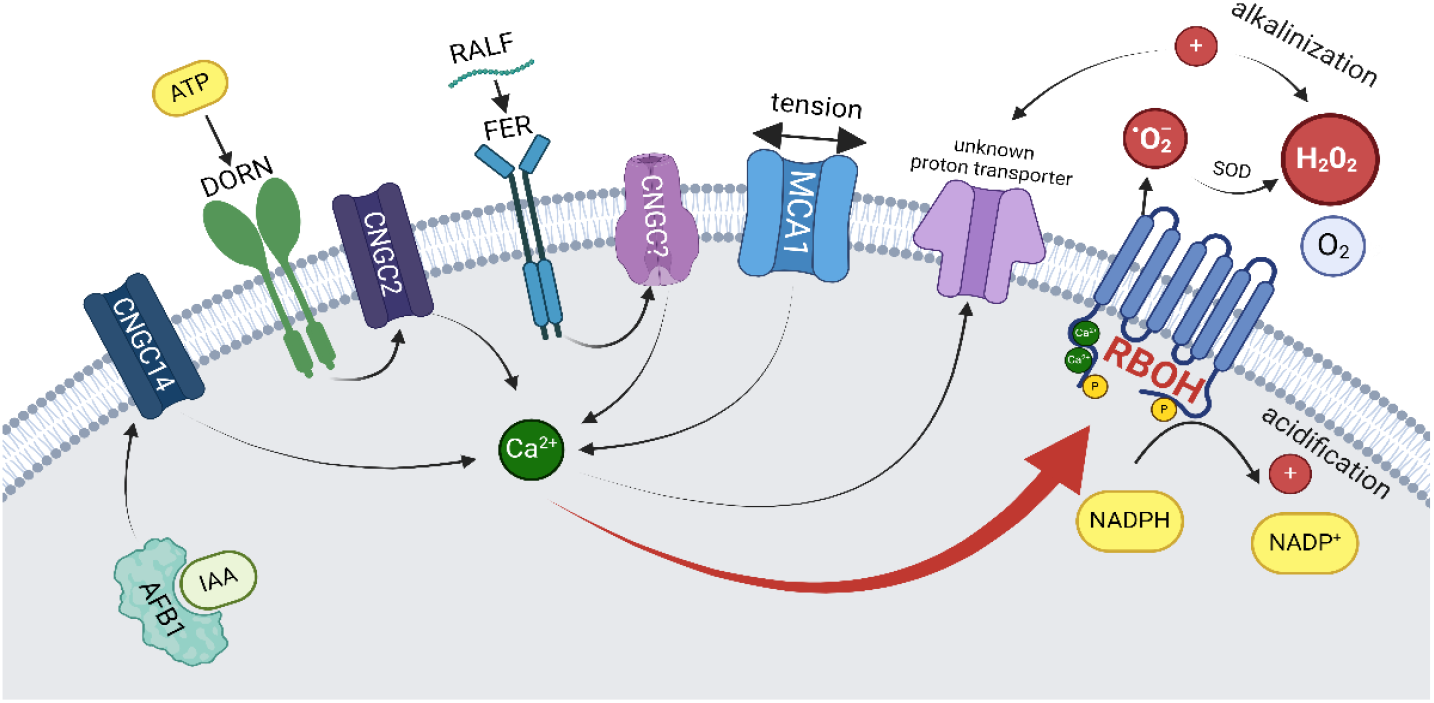
Multiple pathways in root converge on calcium signaling to activate NADPH oxidase. This model is based on the following findings: AFB1 and CNGC14 are required for IAA-induced Ca^2+^ transients (*4, 37*); DORN1 and CNGC2 for eATP-induced Ca^2+^ transients (*17, 41*); Feronia for RALF-induced Ca^2+^ transient depending on an unknown Ca^2+^ channel (*15*). MCA1 induces Ca^2+^ spikes in response to mechanical tension (*48*). NADPH oxidases are activated by the Ca^2+^ transients generated by all these factors (this study) and are solely responsible for the calcium-induced ROS burst (this study). Since IAA induces alkalinization in an RBOH-independent manner, another unknown downstream alkalinization mechanism must exist (this study). Created with BioRender.com.

To confirm that the ROS burst occurs downstream of mechanosensing genetically, we investigated various mutants related to mechanoperception, such as *feronia* (*fer-4* (*14*)), *theseus* (*the1-4* (*46*)) and *mca1* (*47*). We measured the ratio of ROS production on the convex and concave sides of the root following mechanical bending. While *fer-4* and *the1-4* were comparable to WT, *mca1* mutants did not generate any asymmetry in ROS production (convex:concave side ∼1:1), although basal ROS production was maintained (Fig. 3C,D). The MCA-based mechanism acts also during gravitropic bending since *mca1* mutants exhibited the same, faster gravitropic bending as seen with *rbohc/f* (fig. S3A). As MCA1 functions as a stretch-induced Ca^2+^ channel (*48*), our observations put forward the mechanism where MCA-mediated cytosolic Ca^2+^ transients induce NADPH oxidase-dependent apoplastic ROS production, which stiffens the cell wall as shown before (*49*–*51*) and thus limits the bending.

The *mca1* mutant was described to have severe soil impedance phenotypes (*47*), therefore, we also tested the media penetration ability of *rbohc/f* mutants. In our experiment, we observed growth through an air gap to rule out the mechanical influence of the root hairs. While WT plants penetrated the 0.8% agar easily, only a small portion of *rbohc/f* mutants was able to do this, often growing backwards into the air gap (Fig. 3E-G; Movie S5).

In summary, mechanoperception via the MCA1 channel induces Ca^2+^- and NADPH oxidase-dependent apoplastic ROS bursts for cell wall stiffening and soil penetration.

## Discussion

### NADPH oxidases are a major source of apoplastic ROS on the root surface

Apoplastic ROS is a key compound regulating cell wall stiffness and thus growth (*49*–*51*). It can be generated by different enzymes, including NADPH oxidases, class III peroxidases, oxalate oxidases, amine oxidases, and lipoxygenases (*51*). While the inhibitor studies proved inconclusive, the genetic findings revealed that the majority of root apoplastic ROS is generated by RBOH NADPH oxidases, particularly during rapid signaling events and stress responses for growth regulation.

### Multiple signals converge on calcium transients to induce ROS bursts

Apoplastic ROS bursts are triggered by a multitude of signaling pathways, each with a proposed mechanism involving e.g. downstream ROP GTPases (*14*), PtdIns 3-kinase activation (*21*), transcriptional induction of NADPH oxidases (*13*), or their phosphorylation (*52*). The involvement of Ca^2+^ in the activation mechanism of plant NADPH oxidases has been primarily studied in heterologous systems. Research in HEK cells has shown that elevated cytoplasmic Ca^2+^ concentrations are sensed by two N-terminal Ca^2+^-binding EF motifs in RBOHD, triggering conformational changes that lead to enzymatic activation and subsequent ROS production. Another well-established activation mechanism is mediated by phosphorylation (*38, 53*).

Our study contextualizes Ca^2+^ transients within multiple plant signaling pathways upstream of ROS bursts, including those triggered by IAA, eATP, RALF, and mechanical stimuli. All these pathways converge on the induction of cytosolic Ca^2+^ transients, which in turn activate NADPH oxidases. This is evidenced by the absence of ROS bursts in Ca^2+^ channel mutants *cngc14-2* and *mca1-1*, as well as in Ca^2+^-free media. As different sources of Ca^2+^ (CNGCs or MCA1 channels, optogenetic Ca^2+^ channel XXM2.0) are comparably effective, this suggests that Ca^2+^ transients are not only necessary but also sufficient universal signals to induce apoplastic ROS bursts.

### Role of NADPH oxidases and ROS downstream of auxin and mechanical stimulation

The involvement of ROS and NADPH oxidases in root gravitropism has been frequently discussed previously (*12, 54, 55*). Our study clarifies that auxin stimulates ROS bursts via AFB1-CNGC14 signaling module, completely dependent on RBOH NADPH oxidases.

The observation that *rbohc/f* mutants with no detectable auxin-induced ROS burst exhibit normal root growth inhibition and rather enhanced gravitropic bending does not directly disprove the previous findings that asymmetric apoplastic ROS can trigger root bending (*12*). However, it strongly advocates that on the top of the ROS-induced cell wall modulation, auxin triggers also different, more prominent mechanisms of rapid root growth inhibition. This is likely an unknown anion or H^+^ channel-based mechanism, providing means of rapid apoplast alkalinization (*56*). Our results imply that if this mechanism is ever discovered and knocked out, the resulting plants will still retain some degree of gravitropic bending due to auxin-induced NADPH oxidase activity.

In contrast to gravitropic bending, the ROS burst downstream of mechanical receptor and Ca^2+^ channel MCA1 is sufficient and necessary to reinforce the cell wall in the course of media penetration. The mechanism behind the divergence between mechanical and hormonal signaling (both mediated by downstream Ca^2+^) is currently unclear.

### NADPH oxidases and soil impedance response

Our data suggest that the NADPH oxidase-dependent apoplastic ROS burst plays a crucial role in balancing growth in response to hormonal and mechanical signaling, thus aiding root navigation through soil by responding to soil impedance. When a root encounters compacted soil, it bends, the resulting tension on the convex side activates MCA1 mechanical receptor, which generates cytosolic Ca^2+^ transients. Ca^2+^ activates NADPH oxidase to generate apoplastic ROS, which modulates the cell wall. The phenomenon of ROS inducing cell wall rigidity is well established. Various cell wall components, such as hemicelluloses conjugated with phenolic residues, structural proteins, or monolignols, are cross-linked by cell wall peroxidases in an H_2_O_2_-dependent manner, resulting in growth inhibition (*57, 58*). Besides direct cell wall modulation, other processes may be triggered by the plasma membrane ROS receptor HPCA1 (*59*) or by AUX/IAA multimerization (*60*). Previous studies, utilizing cytoplasmic ROS sensors (*10*), which fail to respond to apoplastic ROS bursts, have shown either no change or a decrease in ROS during mechanical bending (*10, 61*).

In summary, from a mechanistic perspective, our results support a model, in which root bending is perceived by MCA1, a mechanosensing Ca^2+^ channel. The resulting cytosolic Ca^2+^ increase activates NADPH oxidases and leads to rapid apoplastic ROS production and alkalinization on the stretched side of the root. This process increases cell wall rigidity, thereby preventing further bending. If this mechanism fails, due to the absence of MCA1 mechanoreceptor or NADPH oxidases, the root would continue bending and ultimately fail to penetrate the soil.

## Author contributions

Conceptualization: IK, JF Methodology: IK, MR, KK, SG, GN Investigation: IK, DV, MR Visualization: IK, MR Funding acquisition: JF Project administration: JF, IK Writing – original draft: IK, JF Writing – review & editing: IK, JF, DV, MR, MC, MW, GN

## Acknowledgments

We gratefully acknowledge the Lab Support Facility (LSF), the Imaging & Optics Facility (IOF) for the support with imaging. We thank to Dr. Matias Fendrych and his team for the help with the microfluidics upgrades.

