## Supplementary material for "Calcium-triggered apoplastic ROS bursts balance gravity and mechanical signals to navigate soil": Methods and Supplementary Figures S1-S4

#### Materials and Methods

#### Materials and methods

##### Plant cultivation

Seed sterilization was executed with chlorine gas or surface sterilization by 96% ethanol. If not stated otherwise, seeds were sown on solid agarose media – half-strength Murashige and Skoog (0.5x MS) medium supplemented with 1% sucrose and 0.8% phyto agar (pH=5.9) – stratified at 4°C for 2 days and subsequently grown vertically at 21°C with 16h light/8h dark cycles. Experiments were performed on the 5-7 day after germination (DAG) old seedlings.

##### Plant treatments

NADPH oxidase inhibitors DPI Diphenyleneiodonium chloride (sigma D2926) and VAS2870 (sigma SML0273) were dissolved DMSO and aliquots were frozen and then added to the media in the indicated concentrations. Corresponding amount of the DMSO was added to the control samples. To induce intracellular $Ca^{2+}$ transients, ATP (Thermo Scientific R0441), IAA (Duchefa I0901) and RALF1 (, At1G02900 A5-49; GL Biochem Lot No. P220237-MX386416) were used from the aliquots stored in water (ATP; 10mM stock, RALF1; 1mM) and 96% ethanol (IAA; 100μM stock). To prepare the $Ca^{2+}$ free MS media, ½ MS was mixed up from the individual salts: Ammonium nitrate (Sigma A3795); Potassium nitrate (Sigma P8291); Magnesium sulfate heptahydrate for analysis (Merck 105886); Potassium phosphate dibasic (Sigma P3786); Manganese sulfate monohydrate (Sigma M7899); Zinc sulphate heptahydrate (sigma Z1001); Potassium iodide (sigma p2963); Copper(II) sulfate pentahydrate (Sigma 209198); Sodium molybdate (sigma 243655); Cobalt(II) chloride (sigma 409332); Boric acid (sigma B6768); Iron(II) sulfate heptahydrate (F8263). These were combined in a 50% amount of the standard MS according to (*1*). Then, calcium chloride dihydrate (Sigma C3306); or Magnesium chloride (Sigma M4880) were added to generate $Ca^{+}$ and $Ca^{-}$ media. In the conditions without $Ca^{2+}$, plants did not respond to IAA with $Ca^{2+}$ spikes, but retained normal growth for first 2 hours after which, they gradually aborted. The experiments were performed within first 20 minutes after the $Ca^{2+}$ depletion.

Mechanical bending was performed as follows: root was placed on the microscopy slide in the droplet of the media. One cover slide was placed at the side of the root, another was used to cover the sample, so that the first cover slide can be gently pushed towards the root during the imaging. Such a setup was mounted to the stage and left for 5 minutes to recover. Then, during the imaging, inner slide was pushed in until visible bending of the root was present (15-30°). The gravitropic bending assays were performed in the vertically oriented scanner where the entire plates were placed, left to rest for 30 minutes and then rotated 90° after which scanning was performed with period of 30 minutes.

##### Transgenic lines and Crispr knockout

All Arabidopsis mutants and transgenic lines which were used in this project are in the Columbia-0 (Col-0) background. Following lines were ordered from the NASC (The European Arabidopsis Stock Centre): *fer-4*, NASC_N69044; *the1-4* nasc_N829966 (*2*); *rbohc*, salk_016593C (*3*). *mca1* mutant (*4*) was kindly donated by (*5*) R2D2 by (*6*) and gCAMP by (*7*). Plants expressing ccvAFB1 were obtained from (*8*). The XXM 2.0 *Arabidopsis* was obtained by transforming the XXM2.0 plasmid from (*9*) into Col-0.

To knock out *RBOHF*, two guide RNAs with no off targets were selected: GCTAATCAAAGCGGCGGTGC; TTAACGGTGATCAAGAGTTC. Using this, two pairs of overlapping primers were designed and used in a PCR reaction with the pCBC-DT1T2 template and then using Golden Gate[tm] reaction into the pHEE401 vector (*10, 11*). Sequenced plasmids were then transformed into the *Arabidopsis thaliana rbohc* mutants (salk_016593C, previously characterized in (*3*)). Candidates selected based on the hygromycin resistance were then subjected to genotyping. Two independent lines which lack the whole 328bp region between the two guide RNAs were selected. In both lines, this results in the protein shortened to I201 followed by 6 frameshifted amino acids and premature stop codon. The CAS9 cassette was then removed by segregation and genotyping using genotyping against hygromycin.

| | |
|---|---|
| RbohFG1_BsF | ATATATGGTCTCGATTGCTAATCAAAGCGGCGGTGCGTT |
| RbohFG1_F0 | TGCTAATCAAAGCGGCGGTGCGTTTTAGAGCTAGAAATAGC |
| RbohFG2_R0 | AACTTAACGGTGATCAAGAGTTCCAATCTCTTAGTCGACTCTAC |
| RbohFG2_RsR | ATTATTGGTCTCGAAACTTAACGGTGATCAAGAGTTCAA |
| Rbohf gen_f | ATGAAACCGTTCTCAAAGAACGATCG |
| Rbohf gen r | CATTGAGCGAAATCGGAGCG |
| Hyg rev | CTTTGCCCTCGGACGAG |
| Hyg_for | ATGAAAAAGCCTGAACTCACCG |

**Microscopy**

For imaging, vertically (Fig. 1-2, fig. S2,3) or horizontally (Fig. 3, fig. S4) mounted Zeiss LSM800 was used with air objective Plan Apochromat 10x/0.45 M27. With either closed 1AU pinhole (Fig. 1, fig. S1-fig. S4) or fully opened pinhole (Fig 2,3, fig. S1C). Microfluidic imaging was performed as introduced in (*12*). Root tracker was used for the supplementary movie 2 (*13*). ROS imaging was performed through the microscopy slide in order to be able to wash the roots. Roots were placed to the edge of the cover glass so that exchange of the media by itself would not trigger $Ca^{2+}$ spike. This effect was negligible as demonstrated in Fig.2B. Samples were left to rest for ~5 minutes, followed by the first was by mock media wash. After this, second media with the elicitor was added, replacing the original media. Media replacement was observed as rapid drop of in the intensity of the Amplex Red signal roughly to the levels of the control Amplex Red sample. DHE (Sigma D7008) imaging was performed using 10µM solution prepared out of 10mM stock solution in DMSO. FS stain (Invitrogen F1130) was prepared in the unbuffered 1/2MS media with sucrose and imaged as described previously (*14*).

**Optogenetics**

The 4 DAG seedling grown under red light were fixed by low melting agarose (Promega, V2111) on the bottom of the glass chamber and the agarose around root tip was cut out. The chamber was then filled with 5ml of ½ MS medium with 1% sucrose and 1µM Amplex Red dye. The imaging was dome on Leica Stellaris LSM with 580nm excitation. And the light treatment was done by external blue LED placed next to objective. For analysis the images were stabilized by StackReg (2.0.0) Fiji (2.15.1) plugin. Amplex Red halo florescence intensity was measured by segmented line tool with thickness 15 around the root tip (from meristem to depreciation zone). Intensity in medium more distant from root was also measured. The signal was normalized to mean value before first treatment and divided be normalized signal from medium outside the halo.

**Data analysis**

Fiji ImageJ bundle was used for the image analysis (*15*). For the ROS measurements, mean pixel value at the time 0 was subtracted from all following values, the data was not normalized. For the mechanical stress (Fig. 3. fig. S4), images were stabilized using Correct 3D Drift plugin of Fiji. For the analysis of the data obtained from the vertical scanner was performed as in (*16*), images were first stabilized using the Correct 3D drift plugin. Angle of each root was measured in each time frame. Relative bending was evaluated, with initial angle set to 0. Root growth ratio for the fig. S2B was calculated from the root increment in 12 hours. Root growth from the microscopy observations was measured as cross-correlation shift of root tip in the direction of the root orientation by chi2_shifts function of image-registration (0.2.9) python package. The root shift was corrected for stage movement by subtracting movement of static background.

For measurement of root bending first the distance from root tip was computed as Travel time transformation by python package scikit-fmm (2023.4.2). Root curvature was calculated as change of distance gradient angle on the root. The mean intensity of GCaMP signal (405/488nm) was calculated in region corresponding to elongation zone in 1st 1/4 of root thickness.

### Fig. S1

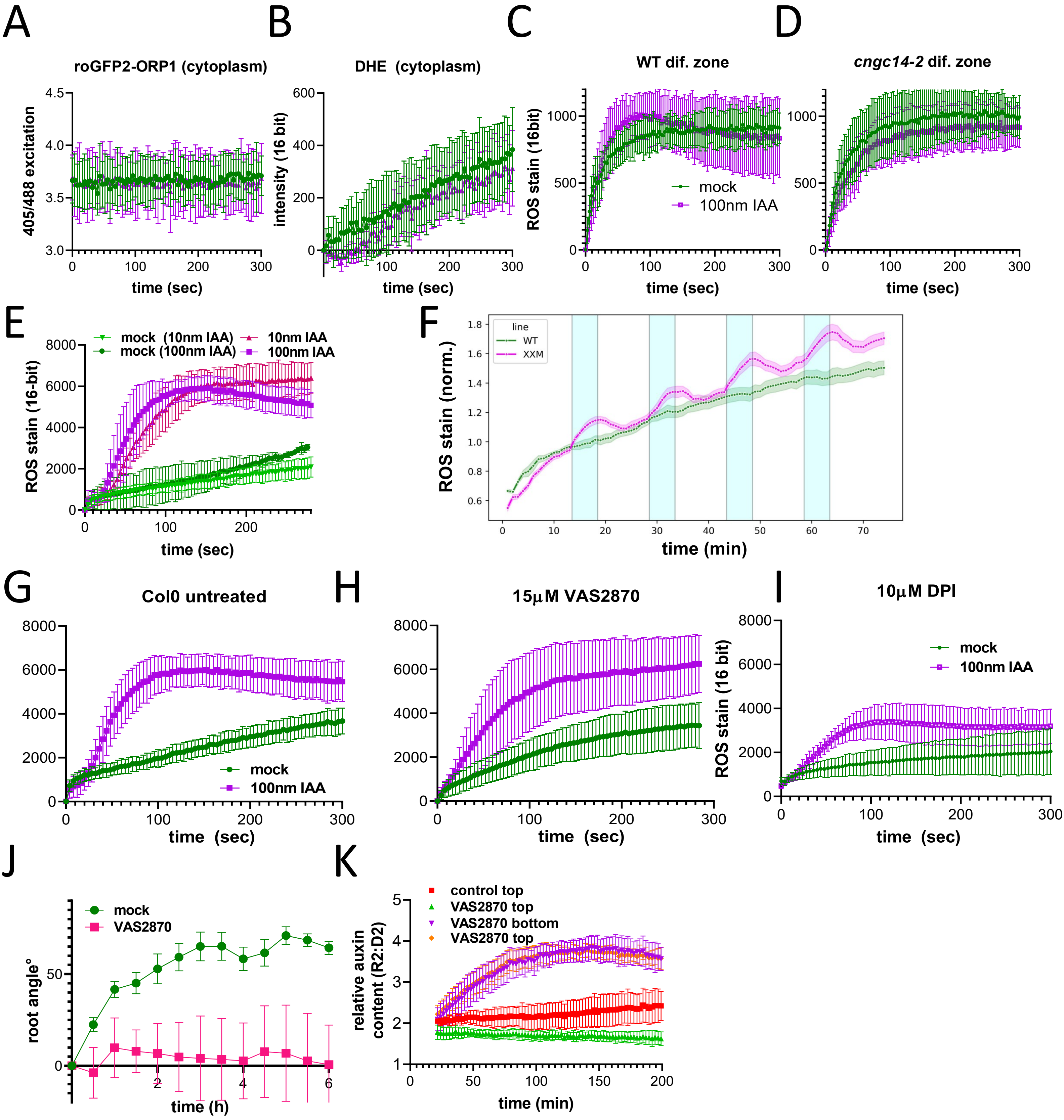
A
roGFP2-ORP1 (cytoplasm)
405/488 excitation
time (sec)
B
DHE (cytoplasm)
intensity (16 bit)
time (sec)
C
WT dif. zone
ROS stain (16bit)
mock
100nm IAA
time (sec)
D
cngc14-2 dif. zone
time (sec)
E
mock (10nm IAA)
mock (100nm IAA)
10nm IAA
100nm IAA
ROS stain (16-bit)
time (sec)
F
line
WT
XXM
ROS stain (norm.)
time (min)
G
Col0 untreated
mock
100nm IAA
time (sec)
H
15μM VAS2870
time (sec)
I
10μM DPI
ROS stain (16 bit)
mock
100nm IAA
time (sec)
J
mock
VAS2870
root angle°
time (h)
K
control top
VAS2870 top
VAS2870 bottom
VAS2870 top
relative auxin content (R2:D2)
time (min)

**Supplementary Figure 1. ROS measurements in the cytoplasm and apoplastic space upon 100 nM IAA and 10 nM IAA induction and VAS2870 impact on root gravitropism**

**(A)** Comparison of cytoplasmic roGFP2-ORP1 ROS detection upon 100 nM IAA treatment and mock treatment.
**(B)** Cytoplasmic ROS accumulation measured as DHE signal intensity upon 100 nM IAA and mock treatment. 10µM DHE was used.
**(C,D)** ROS burst in the root differentiation zone upon 100nm IAA treatment using Amplex Red.
**(E)** Induction of apoplastic ROS by 10 and 100 nM IAA measured using Amplex Red signal intensity.
**(F)** Optogenetic activation of XXM2.0 channel leads to apoplastic ROS burst on the root surface, measured as normalized Amplex Red intensity. Blue stripes represent light treatment. The experiment was repeated 3 times with min. 5 seedlings per line.
**(G-I)** Effect of NADPH oxidase inhibitor VAS2870 and DPI on the 100nm IAA induced ROS burst measured with Amplex Red. 20 min. pretreatment was performed.
**(J)** Root gravitropic bending of control plants and plants treated with 15 µM VAS2870, which resulted in complete loss of the root gravitropism. Plants were transferred to the treatment media and rotated immediately.
**(K)** Loss of the root gravitropism using VAS2870 is accompanied by auxin transport inhibition defects. Auxin distribution detected using the R2D2 reporter during the course of root gravitropic bending. While mock treatment shows higher IAA concentration at the bottom side of the root, 15 µM VAS2870 treatment results in hyperaccumulation of auxin regardless of the gravitropic vector. Imaging and VAS2870 treatments started 20 minutes after transfer of plants and rotation by 90°. See also supplementary movie 2.

Fig. S2

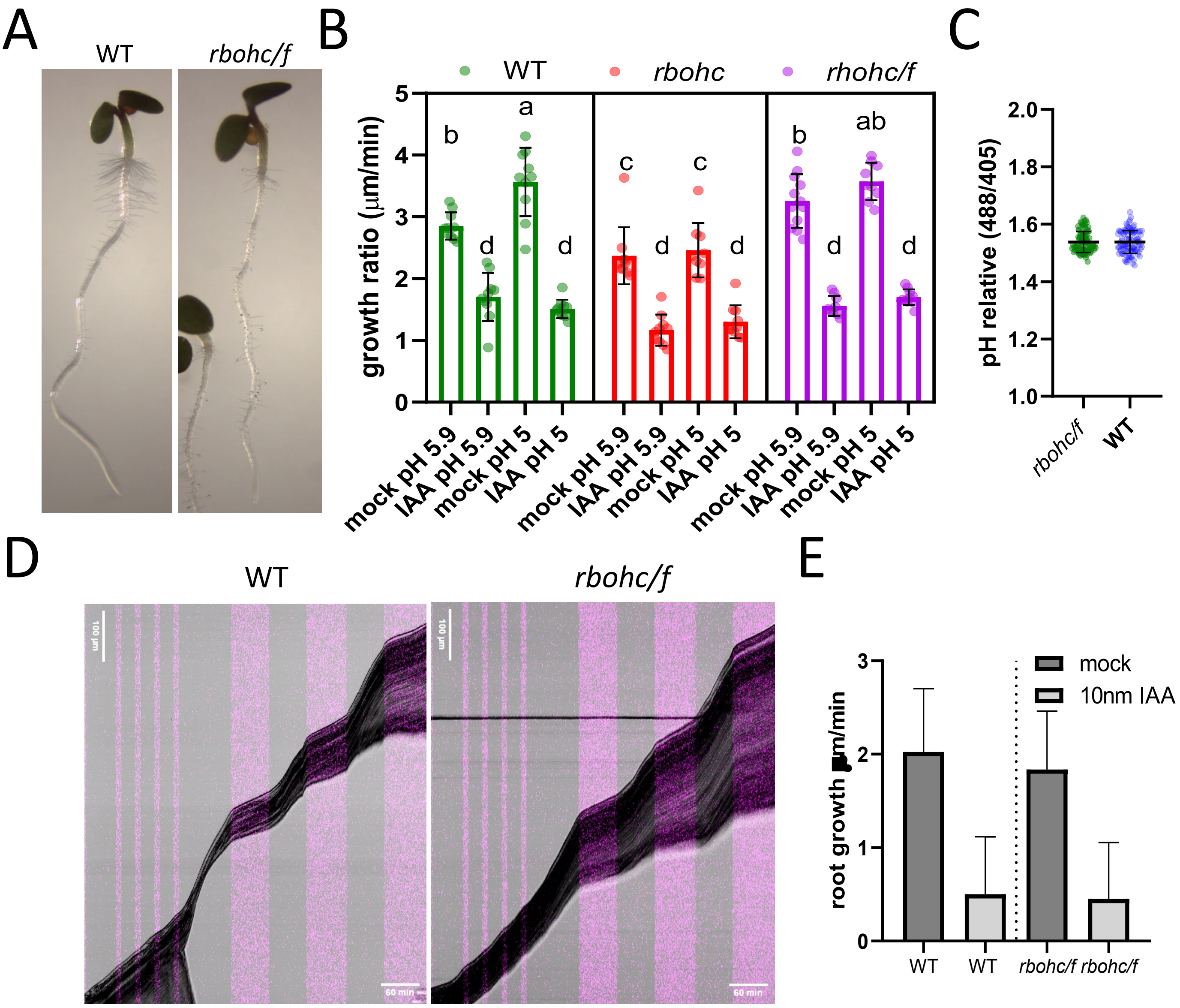


**Supplementary Figure S2. *rbohc/f* double mutants display normal root growth ratio with and without auxin**

**(A)** Representative images of WT and *rbohc/f* double mutants.

**(B)** Long-term effect of 10 nM IAA on the *rbohc* and *rbohc/f* root growth ratio averaged over 30 hours of treatment. Each dot represents an individual root.

**(C)** Overall pH of the *rbohc/f in* the root elongation zone. Five horizontally grown plants were measured on both sides of the root.

**(D)** Kymographs of the microfluidic experiment where roots were treated with a series of 10-minute and 60-minute treatments with 10 nM IAA (magenta stripes). For the video, see Supplementary Movie 3.

**(E)** Quantification of the root growth ratio with and without IAA in the microfluidic setup, averaged over 3 auxin treatments. Similar results were obtained in two additional replicas for 90-100 minutes.

Fig. S3

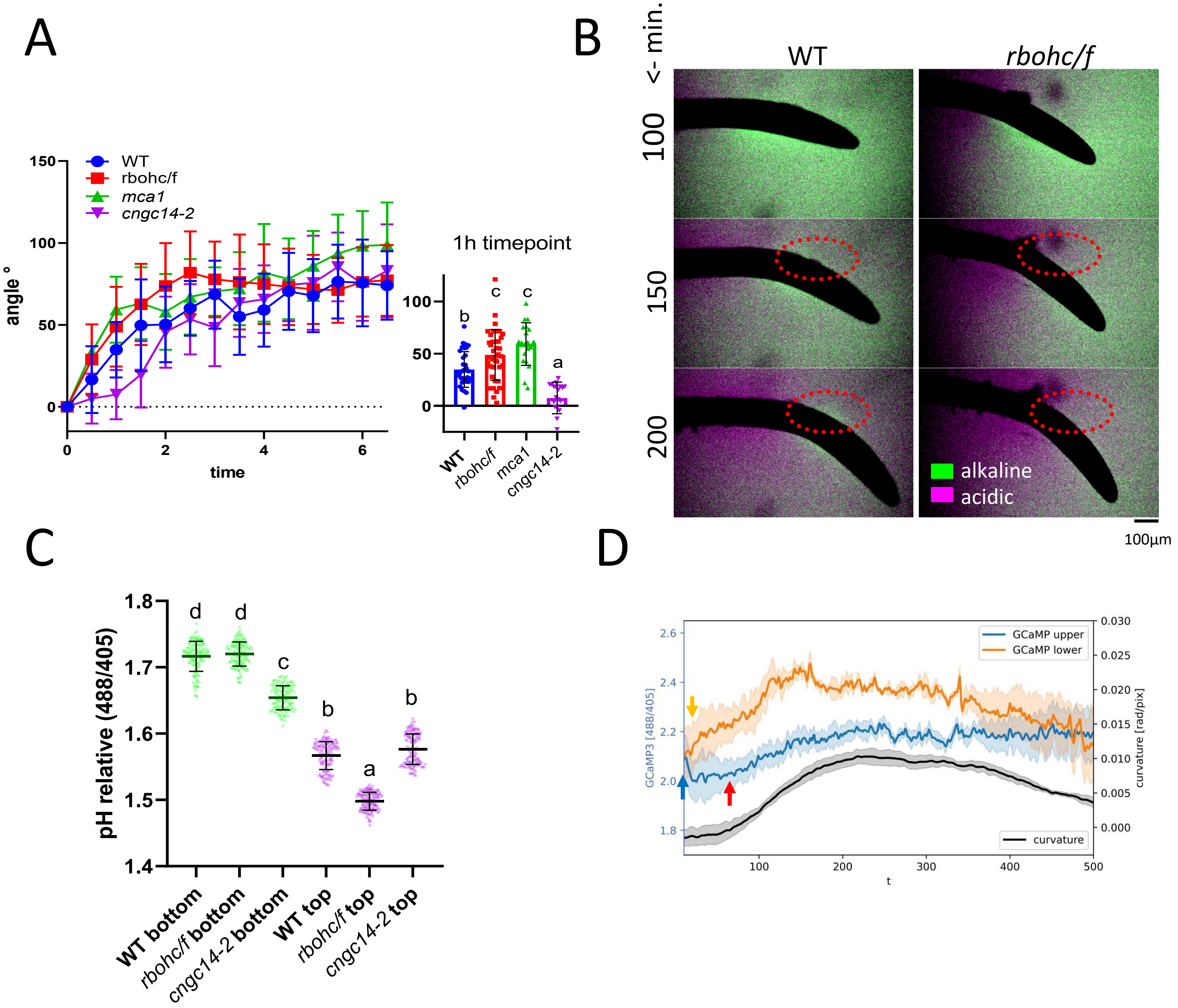


**Supplementary Figure S3. Gravitropic bending of the *rbohc/f* mutants and their relative apoplastic pH in the elongation zone.**

**(A)** Root gravitropic bending of WT, *mca1*, *cngc14-2*, and *rbohc/f* mutants. Inlet depicts timepoint 1h after rotation. Each dot represents one root. Letters denote statistical significance (one-way ANOVA, $p<0.02$)

**(B)** Representative images of the apoplastic pH halo in WT and *rbohc/f* mutants during the course of root gravitropic bending. Red dotted circles highlight the top side of the root with visible alkalinization in WT.

**(C)** Apoplastic pH quantification of the *rbohc/f* and *cngc14-2* mutants at the top and bottom elongation zones 90-160 minutes after rotating the sample 90°. (one-way ANOVA, $p<0.0001$)

**(D)** Calcium profile of WT plants during the course of gravitropic bending imaged immediately after rotating the sample, monitored using the GCaMP 3.0 fluorescent reporter. The blue arrow marks the initial drop in the calcium signal from the top (upper) side of the root associated with statoliths leaving the membrane in the columella, while the orange arrow marks the increase in the calcium signal at the bottom (lower) side after the statoliths touch the new bottom membrane. The red arrow highlights the increase in the calcium signal at the upper side associated with root bending, which appears at the onset of root bending.

### Fig. S4

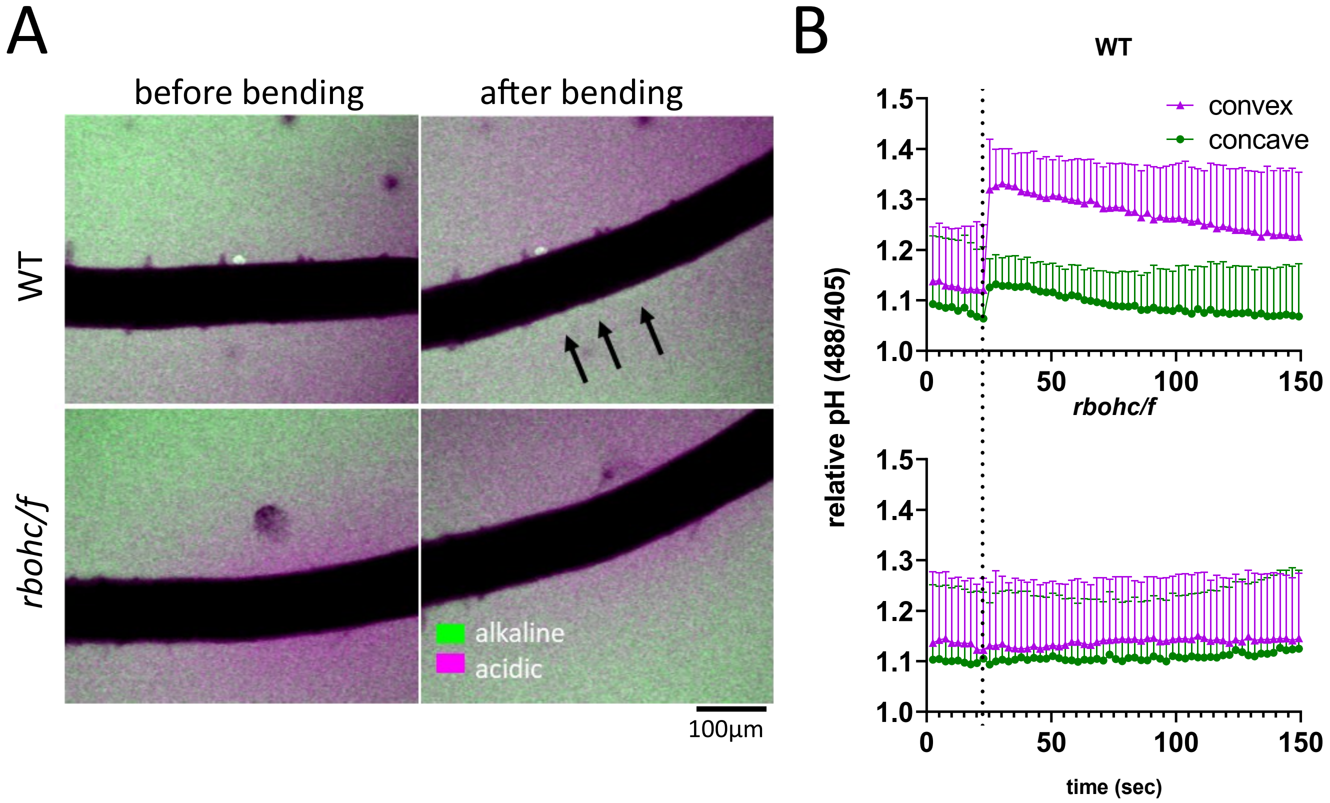


**Supplementary Figure S4. Mechanical stimulus induced alkalinization in the WT and rbohc/f mutants.**

**(A)** Representative videos displaying apoplast alkalinization immediately after root bending (black arrows)
**(B)** quantification of 6 bending experiments with WT and *rbohc/f* mutants. Dotted line depicts the bending event.

**Movie S1:** imaging of $Ca^{2+}$ (green) and ROS (magenta) upon 10nm IAA treatment.

**Movie S2:** 15 µM VAS2870 treatment on R2D2 auxin sensor during the course of root gravitropic bending. Yellow signal depicts DII-VENUS (degraded with rising auxin levels) magenta signal depicts mDII:tdTOMATO signal (ratiometric reference).

**Movie S3:** Fast response of *rbohc/f and WT* to 10nm IAA (magenta)

**Movie S4:** ROS (magenta) upon mechanical bending in WT expressing GCaMP (green) and *rbohc/f*

**Movie S5:** *rbohc/f* agar penetration phenotype
